# PhyKIT: a UNIX shell toolkit for processing and analyzing phylogenomic data

**DOI:** 10.1101/2020.10.27.358143

**Authors:** Jacob L. Steenwyk, Thomas J. Buida, Abigail L. Labella, Yuanning Li, Xing-Xing Shen, Antonis Rokas

**Affiliations:** Vanderbilt University, Department of Biological Sciences, VU Station B #35-1634, Nashville, TN 37235, United States of America; 9 City Place #312, Nashville, TN 37209, United States of America; Ministry of Agriculture Key Lab of Molecular Biology of Crop Pathogens and Insects, Institute of Insect Sciences, Zhejiang University, Hangzhou 310058, China

**Keywords:** molecular phylogenetics, phylogenomics, multiple sequence alignment, phylogenetic signal, network, gene-gene covariation, evolutionary rate, polytomy, tree distance

## Abstract

Diverse disciplines in biology process and analyze multiple sequence alignments (MSAs) and phylogenetic trees to evaluate their information content, infer evolutionary events and processes, and predict gene function. However, automated processing of MSAs and trees remains a challenge due to the lack of a unified toolkit. To fill this gap, we introduce PhyKIT, a toolkit for the UNIX shell environment with 30 functions that process MSAs and trees, including but not limited to estimation of mutation rate, evaluation of sequence composition biases, calculation of the degree of violation of a molecular clock, and collapsing bipartitions (internal branches) with low support. To demonstrate the utility of PhyKIT, we detail three use cases: (1) summarizing information content in MSAs and phylogenetic trees for diagnosing potential biases in sequence or tree data; (2) evaluating gene-gene covariation of evolutionary rates to identify functional relationships, including novel ones, among genes; and (3) identify lack of resolution events or polytomies in phylogenetic trees, which are suggestive of rapid radiation events or lack of data. We anticipate PhyKIT will be useful for processing, examining, and deriving biological meaning from increasingly large phylogenomic datasets. PhyKIT is freely available on GitHub (https://github.com/JLSteenwyk/PhyKIT) and documentation including user tutorials are available online (https://jlsteenwyk.com/PhyKIT).

## Introduction

Multiple sequence alignments (MSAs) and phylogenetic trees are widely used in numerous disciplines, including bioinformatics, evolutionary biology, molecular biology, and structural biology. As a result, the development of user-friendly software that enables biologists to process and analyze MSAs and phylogenetic trees is an active area of research (Kapli *et al*. 2020).

In recent years, numerous methods have proven useful for diagnosing potential biases and inferring biological events in genome-scale phylogenetic (or phylogenomic) datasets. For example, methods that evaluate sequence composition biases in MSAs (Phillips and Penny 2003), signatures of clock-like evolution in phylogenetic trees (Liu *et al*. 2017), phylogenetic treeness (Lanyon 1988; Phillips and Penny 2003), taxa whose long branches may cause variation in their placement on phylogenetic trees (Struck 2014), and others have assisted in summarizing the information content in phylogenomic datasets and improved phylogenetic inference (Felsenstein 1978; Philippe *et al*. 2011; Salichos and Rokas 2013; Doyle *et al*. 2015; Liu *et al*. 2017; Smith *et al*. 2018; Walker *et al*. 2019).

Other methodological innovations include identifying significant gene-gene covariation of evolutionary rate, which has been shown to accurately and sensitively identify genes that have shared functions, are co-expressed, and/or are part of the same multimeric complexes (Sato *et al*. 2005; Clark *et al*. 2012). Furthermore, gene-gene covariation serves as a powerful evolution-based genetic screen for predicting gene function (Brunette *et al*. 2019). Lastly, a recently developed method has enabled the identification of unresolved internal branches or polytomies in species trees (Sayyari and Mirarab 2018; One Thousand Plant Transcriptomes Initiative 2019); such branches can stem from rapid radiation events or from lack of data (Rokas and Carroll 2006).

Despite the wealth of information in MSAs and phylogenetic trees, there is a dearth of tools, especially ones that allow to conduct these analyses in a unified framework. For example, to utilize the functions mentioned in the previous paragraphs, a combination of web-server applications, ‘hard-coded’ scripts available through numerous repositories and supplementary material, standalone software, and/or extensive programming in languages including R, Python, or C is currently required (Cock *et al*. 2009; Junier and Zdobnov 2010; Revell 2012; Talevich *et al*. 2012; Struck 2014; Kück and Longo 2014; Wolfe and Clark 2015; One Thousand Plant Transcriptomes Initiative 2019). As a result, integrating these functions into bioinformatic pipelines is challenging, reducing their accessibility to the scientific community.

To facilitate the integration of these methods into bioinformatic pipelines, we introduce PhyKIT, a UNIX shell toolkit with 30 functions (Table 1) with broad utility for analyzing and processing MSAs and phylogenetic trees. Current functions implemented in PhyKIT include measuring topological similarity of phylogenetic trees, creating codon-based MSAs, concatenating sets of MSAs into phylogenomic datasets, editing and/or viewing alignments and phylogenetic trees, and identifying putatively spurious homologs in MSAs. We highlight three uses of PhyKIT: (1) calculating diverse statistics that summarize the information content and potential biases (e.g., sequence- or phylogeny-based biases) in MSAs and phylogenetic trees; (2) creating a gene-gene covariation network of evolutionary rates; and (3) inferring the presence of polytomies from phylogenomic data. The diverse functions implemented in PhyKIT will likely be of interest to bioinformaticians, molecular biologists, evolutionary biologists, and others.

**Table 1.**
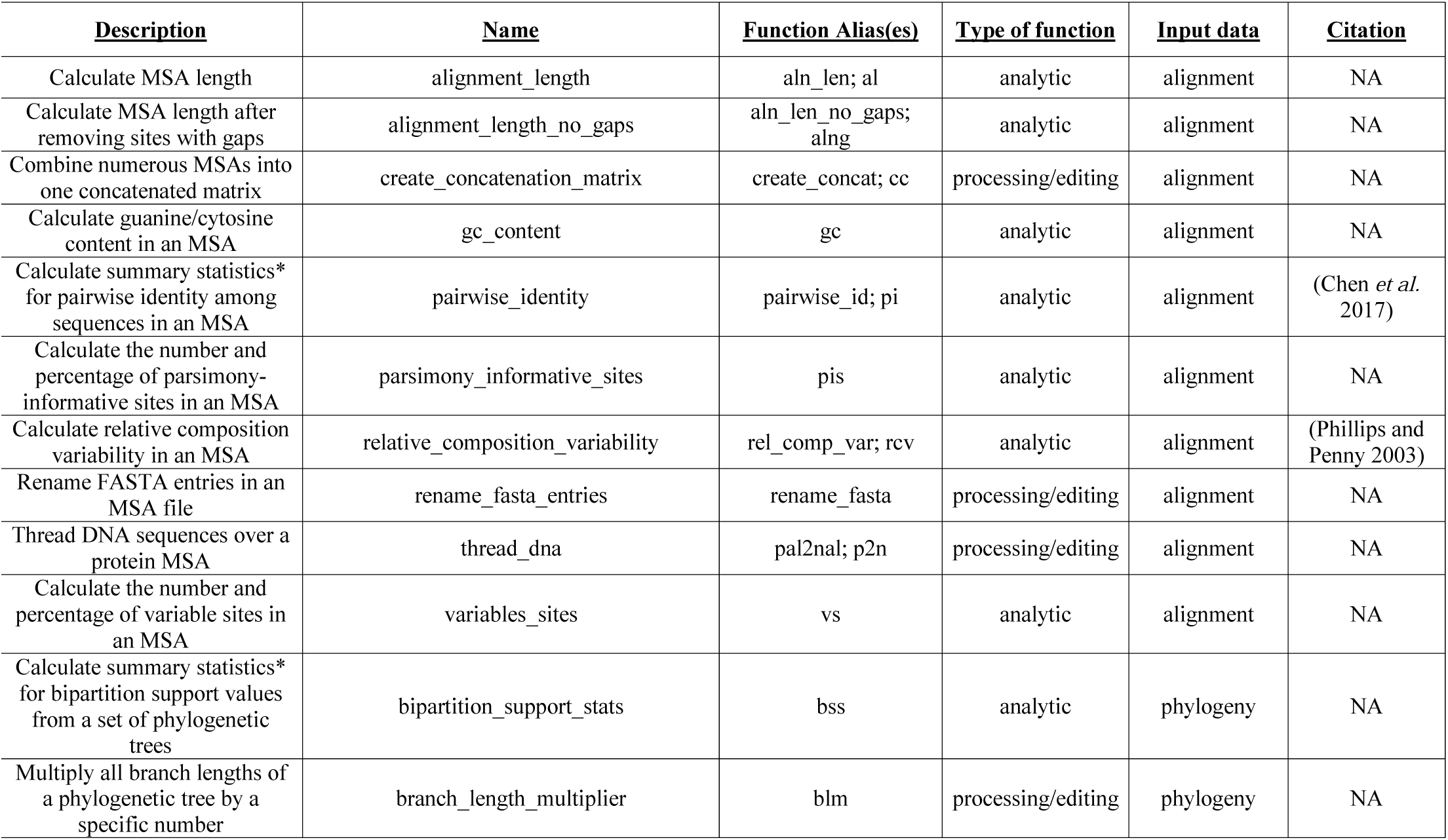
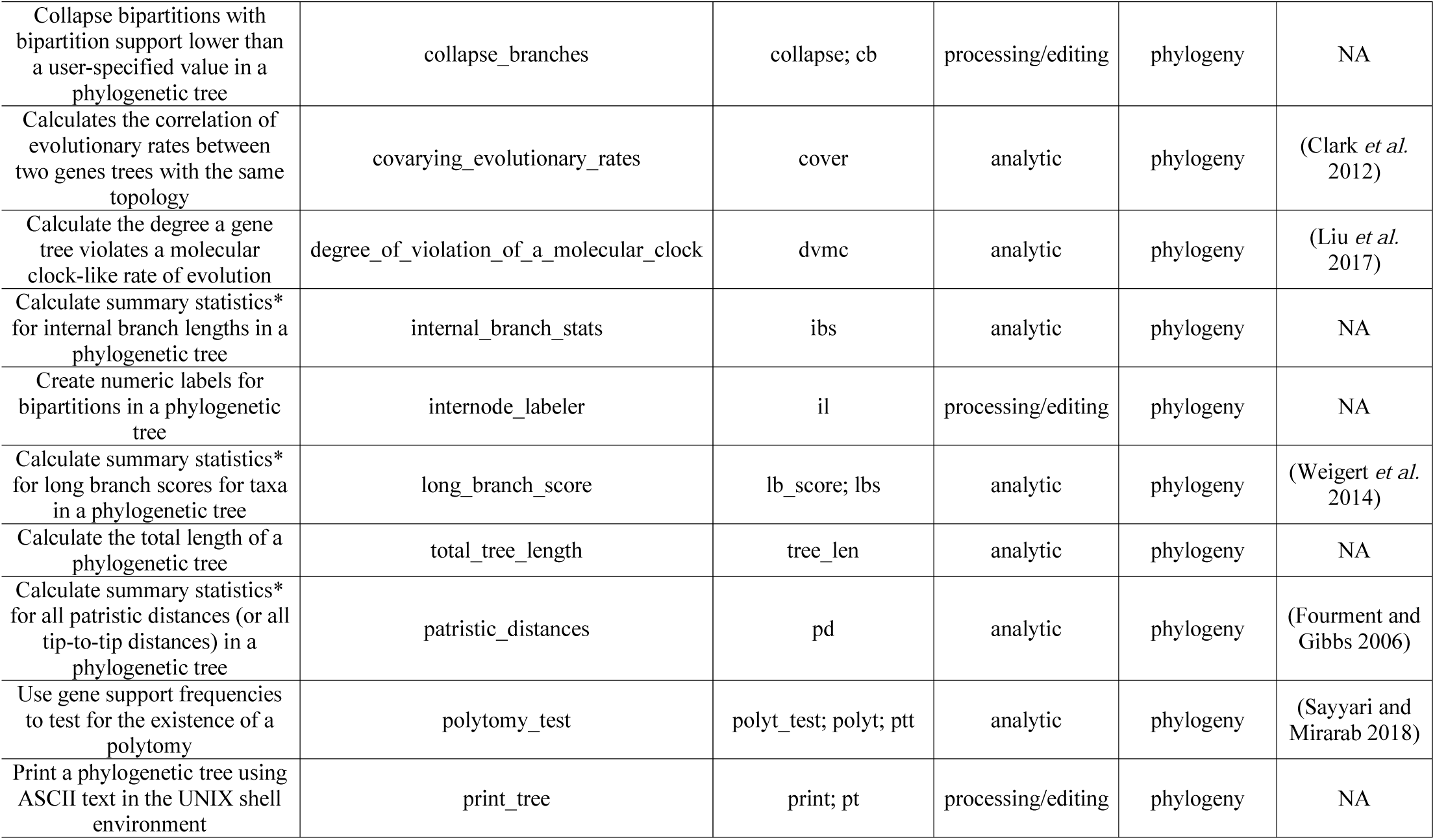
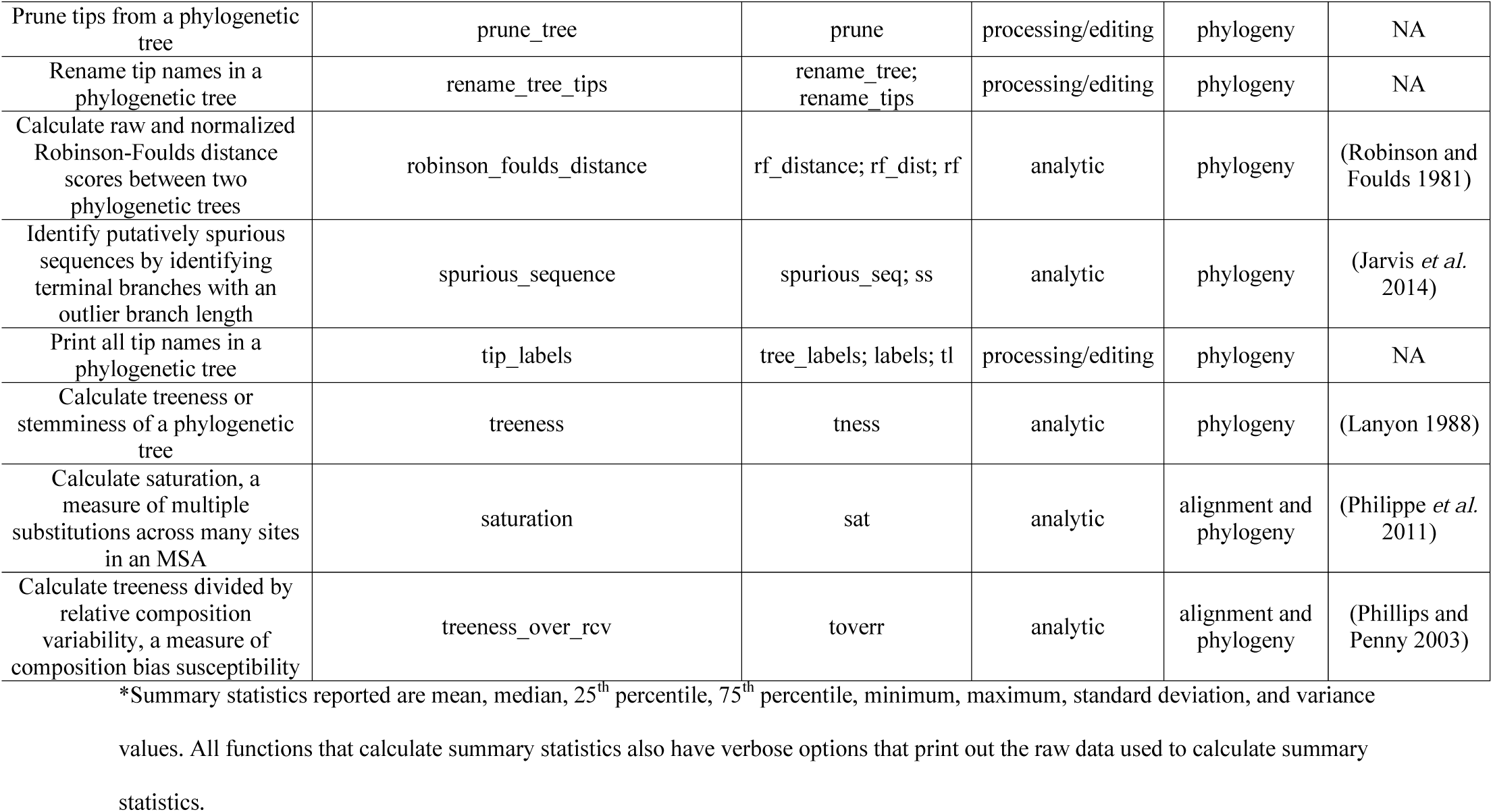
Summary of 30 functions implemented in PhyKIT

## Materials and Methods

PhyKIT is a command line tool for the UNIX shell environment written in the Python programming language (https://www.python.org/). PhyKIT requires few dependencies (Biopython (Cock *et al*. 2009) and SciPy (Virtanen *et al*. 2020)) making it user-friendly to install and integrate into existing bioinformatic pipelines. Furthermore, the online documentation of PhyKIT comes complete with tutorials that detail how to use various functions. Lastly, PhyKIT is modularly designed to allow straightforward integration of additional functions in future versions.

PhyKIT has 30 different functions that help process and analyze MSAs and phylogenetic trees (Table 1). The 30 functions can be grouped into broad categories that assist in conducting analyses of MSAs and phylogenies or in processing/editing them. For example, “analysis” functions help examine information content biases, gene-gene covariation, and polytomies in phylogenomic datasets; “processing/editing” functions help prune tips from phylogenies, collapse poorly supported bipartitions in phylogenetic trees, concatenate sets of MSAs into a single data matrix, or create codon-based alignments from protein alignments and their corresponding nucleotide sequences.

Detailed information about each one of PhyKIT’s functions and tutorials for using the software can be found in the online documentation (https://jlsteenwyk.com/PhyKIT). Here, we focus on three specific groups of functions implemented in PhyKIT that enable researchers to summarize information content in phylogenomic datasets, create gene-gene evolutionary rate covariation networks, and identifying polytomies in phylogenomic data.

### Evaluating information content and biases in phylogenomic datasets

MSAs and phylogenetic trees are frequently examined to evaluate their information content and potential biases in characteristics such as sequence composition or branch lengths (Phillips and Penny 2003; Philippe *et al*. 2011; Struck 2014; Doyle *et al*. 2015; Shen *et al*. 2016a; Liu *et al*. 2017; Smith *et al*. 2018). PhyKIT implements numerous functions for doing so. Here, we demonstrate the application of 14 functions:

#### (1) Alignment length

The length of a multiple sequence alignment, which is associated with robust bipartition support and tree accuracy (Shen *et al*. 2016a; Walker *et al*. 2019);

#### (2) Alignment length with no gaps

The length of a multiple sequence alignment after excluding sites with gaps, which is associated with robust bipartition support and tree accuracy (Shen *et al*. 2016a);

#### (3) Degree of violation of a molecular clock (DVMC)

A metric used to determine the clock-like evolution of a gene using the standard deviation of branch lengths for a single gene tree (Liu *et al*. 2017). DVMC is calculated using the following formula:

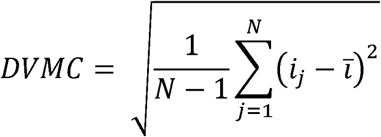

where *N* represents the number of tips in a phylogenetic tree, *i*_*j*_ being the distance between the root of the tree and species *j*, and 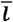represents the average root to tip distance. DVMC can be used to identify genes with clock-like evolution for divergence time estimation (Liu *et al*. 2017);

#### (4) Internal branch lengths

Summary statistics of internal branch lengths in a phylogenetic tree are reported including mean, median, 25^th^ percentile, 75^th^ percentile, minimum, maximum, standard deviation, and variance values. Examination of internal branch lengths is useful in evaluating phylogenetic tree shape;

#### (5) Long branch score

A metric that examines the degree of taxon-specific long branch attraction (Struck 2014; Weigert *et al*. 2014). Long branch scores of individual taxa are calculated using the following formula:

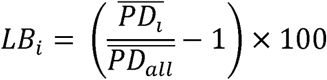

where 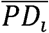 represents the average pairwise patristic distance of taxon *i* to all other taxa, 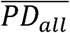 represents the average patristic distance across all taxa, and *LB*_*i*_ represents the long branch score of taxon *i*. Long branch scores can be used to evaluate heterogeneity in tip-to-root distances and identify taxa that may be susceptible to long branch attraction;

#### (6) Pairwise identity

Pairwise identity is a crude approximation of the evolutionary rate of a gene and is calculated by determining the average number of sites in an MSA that are the same character state between all pairwise combinations of taxa. This can be used to group genes based on their evolutionary rates (e.g., faster-evolving genes vs. slower-evolving ones) (Chen *et al*. 2017);

#### (7) Patristic distances

Patristic distances refer to all distances between all pairwise combinations of tips in a phylogenetic tree (Fourment and Gibbs 2006), which can be used to evaluate the rate of evolution in gene trees or taxon sampling density in species trees;

#### (8) Parsimony-informative sites

Parsimony-informative sites are those sites in an MSA that have a least two character states (excluding gaps) that occur at least twice (Kumar *et al*. 2016); the number of parsimony-informative sites is associated with robust bipartition support and tree accuracy (Shen *et al*. 2016a; Steenwyk *et al*. 2020);

#### (9) Variable sites

Variable sites are those sites in an MSA that contain at least two different character states (excluding gaps) (Kumar *et al*. 2016); the number of variable sites is associated with robust bipartition support and tree accuracy (Shen *et al*. 2016a);

#### (10) Relative composition variability

Relative composition variability is the average variability in the sequence composition among taxa in an MSA. Relative composition variability is calculated using the following formula:

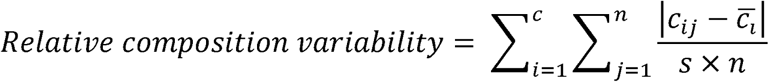

where *c* is the number of different character states per sequence type, *n* is the number of taxa in an MSA, *c*_*ij*_ is the number of occurrences of the *i*th character state for the *j*th taxon, 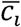 is the average number of the *i*th *c* character state across *n* taxa, and *s* refers to the total number of sites (characters) in an MSA. Relative composition variability can be used to evaluate potential sequence composition biases in MSAs, which in turn violate assumptions of site composition homogeneity in standard models of sequence evolution (Phillips and Penny 2003);

#### (11) Saturation

Saturation refers to when an MSA contains many sites that have experienced multiple substitutions in individual taxa. Saturation is estimated from the slope of the regression line between patristic distances and pairwise identities. Saturated MSAs have reduced phylogenetic information and can result in issues of long branch attraction (Lake 1991; Philippe *et al*. 2011);

#### (12) Total tree length

Total tree length refers to the sum of internal and terminal branch lengths and is calculated using the following formula:

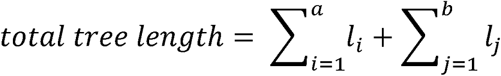

Where *l*_*i*_ is the branch length of the *i*th branch of *a* internal branches and *l*_*j*_ is the branch length of the *j*th branch of *b* terminal branches. Total tree length measures the inferred total amount or rate of evolutionary change in a phylogenetic tree;

#### (13) Treeness

Treeness (also referred to as stemminess) is a measure of the inferred relative amount or rate of evolutionary change that has taken place on internal branches of a phylogenetic tree (Lanyon 1988; Phillips and Penny 2003) and is calculated using the following formula:

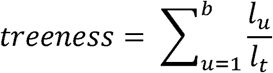

where *l*_*u*_ is the branch length of the *u*th branch of *b* internal branches, and *l*_*t*_ refers to the total branch length of the phylogenetic tree. Treeness can be used to evaluate how much of the total tree length is observed among internal branches;

#### (14) Treeness divided by relative composition variability

This function combines two metrics to measure both composition bias and other biases that may negatively influence phylogenetic inference. High treeness divided by relative composition variability values have been shown to be less susceptible to sequence composition biases and are associated with robust bipartition support and tree accuracy (Phillips and Penny 2003; Shen *et al*. 2016a).

### Calculating gene-gene evolutionary rate covariation or coevolution

Genes that share similar rates of evolution through speciation events (or coevolve) tend to have similar functions, expression levels, or are parts of the same multimeric complexes (Sato *et al*. 2005; Clark *et al*. 2012). Thus, identifying significant coevolution between genes (i.e., identifying genes that are significantly correlated in their evolutionary rates across speciation events) can be a powerful evolution-based screen to determine gene function (Brunette *et al*. 2019).

To measure gene-gene evolutionary rate covariation, PhyKIT implements the mirror tree method (Pazos and Valencia 2001; Sato *et al*. 2005), which examines whether two trees have correlated branch lengths. Specifically, PhyKIT calculates the Pearson correlation coefficient between branch lengths in two phylogenetic trees that share the same tips and topology. To account for differences in taxon representation between the two trees, PhyKIT first automatically determines which taxa are shared and prunes one or both such that the same set of taxa is present in both trees. PhyKIT requires that the two input trees have the same topology, which is typically the species tree topology inferred from whole genome or proteome data. Thus, the user will typically first estimate a gene’s branch lengths by constraining the topology to match that of the species tree. When running this function, users should be aware that many biological factors, such as horizontal transfer (Doolittle and Bapteste 2007), incomplete lineage sorting (Degnan and Salter 2005), and introgression / hybridization (Sang and Zhong 2000), can lead to gene histories that deviate from the species tree. In these cases, constraining a gene’s history to match that of a species may lead to errors in the covariation analysis.

Due to factors including time since speciation and mutation rate, correlations between uncorrected branch lengths result in a high frequency of false positive correlations (Sato *et al*. 2005; Clark *et al*. 2012; Chikina *et al*. 2016). To ameliorate the influence of these factors, PhyKIT first transforms branch lengths into relative rates. To do so, branch lengths are corrected by dividing the branch length in the gene tree by the corresponding branch length in the species tree. Previous work revealed that one or a few outlier branch length values can be responsible for false positive correlations and should be removed prior to analysis (Clark *et al*. 2012). Thus, PhyKIT removes outlier data points defined as having corrected branch lengths greater than five (i.e., removing gene tree branch lengths that are five or more times greater than their corresponding species tree branch lengths). Lastly, values are converted into relative rates using a Z-transformation. The resulting relative rates are used when calculating Pearson correlation coefficients.

### Identifying polytomies in phylogenomic data

Rapid radiations or diversification events have occurred throughout the tree of life including among mammals, birds, plants, and fungi (Jarvis *et al*. 2014; Liu *et al*. 2017; One Thousand Plant Transcriptomes Initiative 2019; Li *et al*. 2020). Polytomies correspond to internal branches whose length is 0 (or statistically indistinguishable from 0) and can be driven either by biological (e.g., rapid radiations) or analytical (e.g., low amount of data) factors. Thus, polytomies are useful for inferring rapid radiation or diversification events and exploring incongruence in phylogenies (Sayyari and Mirarab 2018; One Thousand Plant Transcriptomes Initiative 2019; Li *et al*. 2020).

To identify polytomies, a modified approach to a previous strategy was implemented (Sayyari and Mirarab 2018). More specifically, the support for three alternative topologies is calculated among all gene trees from a phylogenomic dataset. For example, in species tree *((A,B),C), D);*, if examining the presence of a polytomy at the ancestral bipartition of tips *A, B*, and *C*, PhyKIT will determine the number of gene trees that support *((A,B),C);, ((A,C),B);*, and *((B,C),A);* using the rooted gene trees provided by the user. Equal support for the three topologies (i.e., the presence of a polytomy) among a set of gene trees is assessed using a Chi-squared test. Failing to reject the null hypothesis is indicative of a polytomy (Sayyari and Mirarab 2018). Note that this approach is distinct from the approach of Sayyari and Mirarab to identify polytomies because PhyKIT uses a gene-based signal rather than a quartet-based signal. The difference between the two methods is that each gene contributes equally to the inference of a polytomy when a gene-based signal is used, whereas genes with greater taxon representation (which contain a greater number of quartets) will contribute a greater signal during polytomy identification when a quartet-based signal is used.

## Results and Discussion

We outline three example uses of PhyKIT: 1) summarizing information content and identifying potential biases in animal, plant, yeast, and filamentous fungal phylogenomic datasets (Shen *et al*. 2016b; Steenwyk *et al*. 2019; Laumer *et al*. 2019; One Thousand Plant Transcriptomes Initiative 2019), 2) constructing a network of significant gene-gene covariation, which reveals genes of shared functions from empirical data spanning ∼550 million years of evolution among fungi (Shen *et al*. 2020), and 3) illustrating how to identify polytomies using simulated and empirical data (Steenwyk *et al*. 2019).

### Summarizing information content and biases in phylogenomic data

Examining information content in phylogenomic datasets can help diagnose potential biases that stem from low signal-to-noise ratios, multiple substitutions, non-clocklike evolution, and other biological or analytical factors. To demonstrate the utility of PhyKIT to summarize the information content in phylogenomic datasets, we calculated 14 different metrics known to help diagnose potential biases in phylogenomic datasets or be associated with accurate and well supported phylogenetic inferences (Felsenstein 1978; Phillips and Penny 2003; Philippe *et al*. 2011; Struck 2014; Doyle *et al*. 2015; Shen *et al*. 2016a; Liu *et al*. 2017; Smith *et al*. 2018) using four empirical phylogenomic datasets from animals (201 tips; 2,891 genes) (Laumer *et al*. 2019), budding yeast (332 taxa; 2,408 genes) (Shen *et al*. 2018), filamentous fungi (93 taxa; 1,668 genes) (Steenwyk *et al*. 2019), and plants (1,124 taxa; 403 genes) (One Thousand Plant Transcriptomes Initiative 2019) (Figure 1, Table 1).

**Figure 1.**
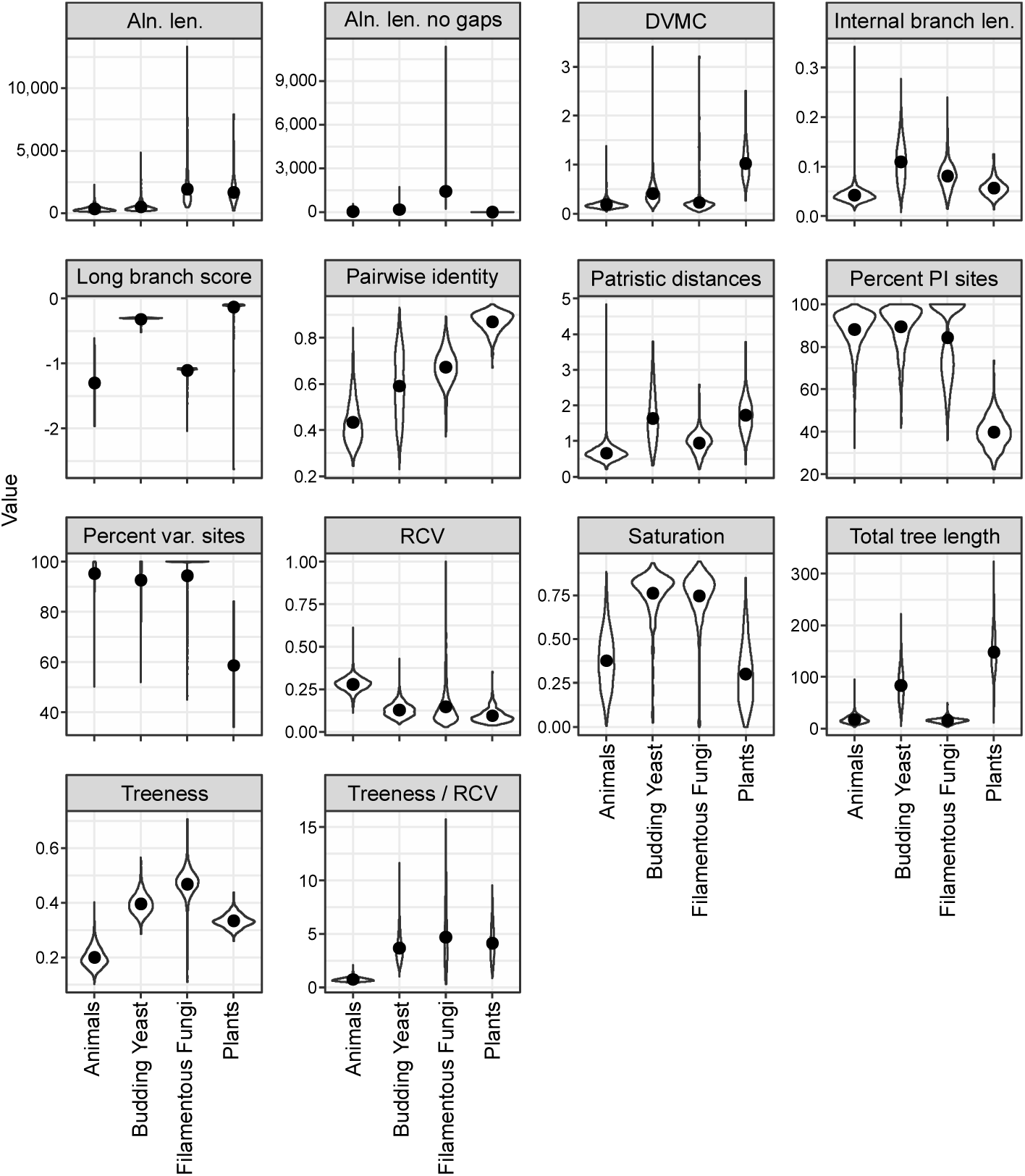
Summary of information content in four empirical phylogenomic datasets. 14 metrics implemented in PhyKIT help summarize the information content and identify potential biases in phylogenomic datasets. Each graph displays a violin plot with a black point representing the mean. Error bars indicate one standard error above and below the mean; however, these are difficult to see in nearly all graphs because they were often near the mean. Abbreviations are as follows: Aln. len.: alignment length; Aln. len. no gaps: alignment length excluding sites with gaps; DVMC: degree of violation of a molecular clock; Internal branch len.: average internal branch length; Patristic distances: average patristic distance in a gene tree; Percent PI Sites: percentage of parsimony-informative sites in an MSA; Percent var. sites: percentage of variable sites in an MSA; RCV: relative composition variability.

Examination of the distributions of the values of the 14 different metrics revealed inter- and intra-dataset heterogeneity (Figure 1). For example, inter-dataset heterogeneity was observed among animal and plant datasets, which had the lowest and highest average pairwise identity across alignments, respectively; intra-dataset heterogeneity was observed in the uniform distribution of pairwise identities in the budding yeast datasets. Similarly, inter-dataset heterogeneity was observed in estimates of saturation where the budding yeast and filamentous fungal MSAs were less saturated by multiple substitutions than the plant and animal datasets; intra-data heterogeneity was also observed in all four datasets. Varying degrees of inter- and intra-dataset heterogeneity was observed for other information content statistics, which may be due biological (e.g., mutation rate) or analytical factors (e.g., taxon sampling, distinct alignment, trimming, and tree inference strategies).

In summary, PhyKIT is useful for examining the information content of phylogenomic datasets. For example, the generation of different phylogenomic data submatrices by selecting subsets of genes or taxa with certain properties (e.g., retention of genes with the highest numbers of parsimony-informative sites or following removal of taxa with high long branch scores) can facilitate the exploration of the robustness of species tree inference or estimating time since divergence (Salichos and Rokas 2013; Liu *et al*. 2017; Shen *et al*. 2018, 2020; Steenwyk *et al*. 2019; Walker *et al*. 2019; Li *et al*. 2020).

### A network of gene-gene covariation reveals neighborhoods of genes with shared function

Genes with similar evolutionary histories often have shared functions, are co-expressed, or are parts of the same multimeric complexes (Sato *et al*. 2005; Clark *et al*. 2012). Using PhyKIT, we examined gene-gene covariation using 815 genes spanning 1,107 genomes and ∼563 million years of evolution among fungi (Shen *et al*. 2020). By examining 331,705 pairwise combinations of genes, we found 298 strong signatures of gene-gene covariation (defined as r > 0.825). The two genes with the strongest signatures of covariation were *SEC7* and *TAO3* (r = 0.87), suggesting that their protein products have similar or shared functions. Supporting this hypothesis, Sec7p contributes to cell-surface growth in the model yeast *Saccharomyces cerevisiae* (Novick and Schekman 1979) and genes with the Sec7 domain are transcriptionally coregulated with yeast-hyphal switches in the human pathogen *C. albicans* (Song *et al*. 2008). Similarly, Tao3p in both *S. cerevisiae* and *C. albicans* is part of a RAM signaling network, which controls hyphal morphogenesis, polarized growth, and cell-cycle related processes including cell separation, cell proliferation, and phase transitions (Bogomolnaya *et al*. 2006; Song *et al*. 2008).

Complex relationships of gene-gene covariation can be visualized as a network (Figure 2). Examination of network neighborhoods identified groups of genes that have shared functions and are parts of the same multimeric complexes. For example, the proteins encoded by *NDC80* and *NUF2* are part of the same kinetochore-associated complex termed the NDC80 complex—which is required for efficient mitosis (Sundin *et al*. 2011)—and significantly covary with one another (r = 0.84). Similarly, multiple genes that encode proteins involved in DNA replication and repair (i.e., *POL2, MSH6, RAD26, CDC9*, and *EXO1*) were part of the same network neighborhood, consistent with previous work suggesting an intimate interplay between DNA replication and multiple DNA repair pathways (Tsubouchi and Ogawa 2000; Lujan *et al*. 2012; Boiteux and Jinks-Robertson 2013). Similarly, network neighborhoods of genes involved in ribosome biogenesis, Golgi apparatus-related transport, and control of DNA replication were identified (Figure 2).

**Figure 2.**
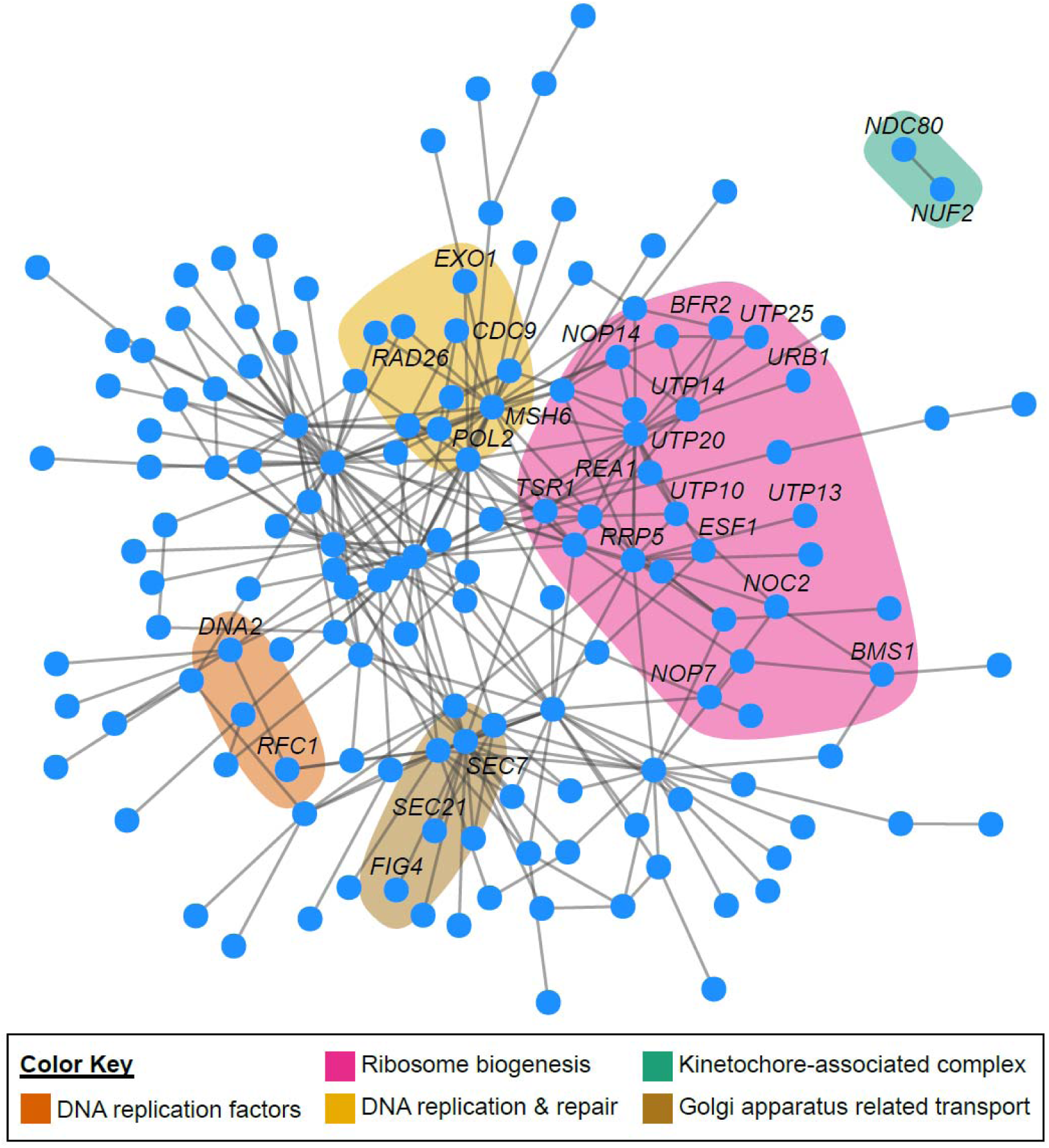
Gene-gene covariation network inferred from ∼550 million years of evolution across 1,107 fungi. A network of significant gene-gene coevolution identifies network neighborhoods representative of associated functional categories. For example, the *NDC80* and *NUF2* genes (toward the top right of the network) were identified to be significantly coevolving with one another (r = 0.84, p < 0.01, Pearson’s correlation test); they both encode proteins that are part of the same multimeric kinetochore-associated complex (green). Similarly, genes that are DNA replication factors (orange), contribute to DNA replication and repair processes (yellow), participate in Golgi apparatus-related transport (brown), or ribosome biogenesis (pink) were found to be neighbors in the network. Network visualization was done with the igraph package, v1.2.4.2 (Hunter and Cohen 2007), in R, v3.6.2 (https://www.r-project.org/).

Taken together, these results indicate PhyKIT is a useful tool for evaluating gene-gene covariation and predicting genes’ functions (Sato *et al*. 2005; Clark *et al*. 2012; Brunette *et al*. 2019). Thus, we anticipate PhyKIT will be helpful for evaluating gene-gene covariation and conducting evolution-based screens for gene functions across the tree of life.

### Identifying polytomies in phylogenomic datasets

Rapid radiations or diversification events have occurred throughout the tree of life (Jarvis *et al*. 2014; Liu *et al*. 2017; One Thousand Plant Transcriptomes Initiative 2019; Li *et al*. 2020). One approach to identifying rapid radiations is by testing for the existence of polytomies in species trees (Sayyari and Mirarab 2018; One Thousand Plant Transcriptomes Initiative 2019; Li *et al*. 2020). Polytomies can also arise when the amount of data at hand is insufficient for resolution (Walsh *et al*. 1999). To demonstrate the utility of PhyKIT to identify polytomies, we tested it using a simulated set of phylogenies that had a branch whose length was extremely small (Figure 3A). We found that PhyKIT was able to conservatively identify the simulated polytomy. Our results demonstrate PhyKIT can accurately identify a polytomy and provide further support that equal support among alternative topologies can be used as a means to identify a rapid radiation.

**Figure 3.**
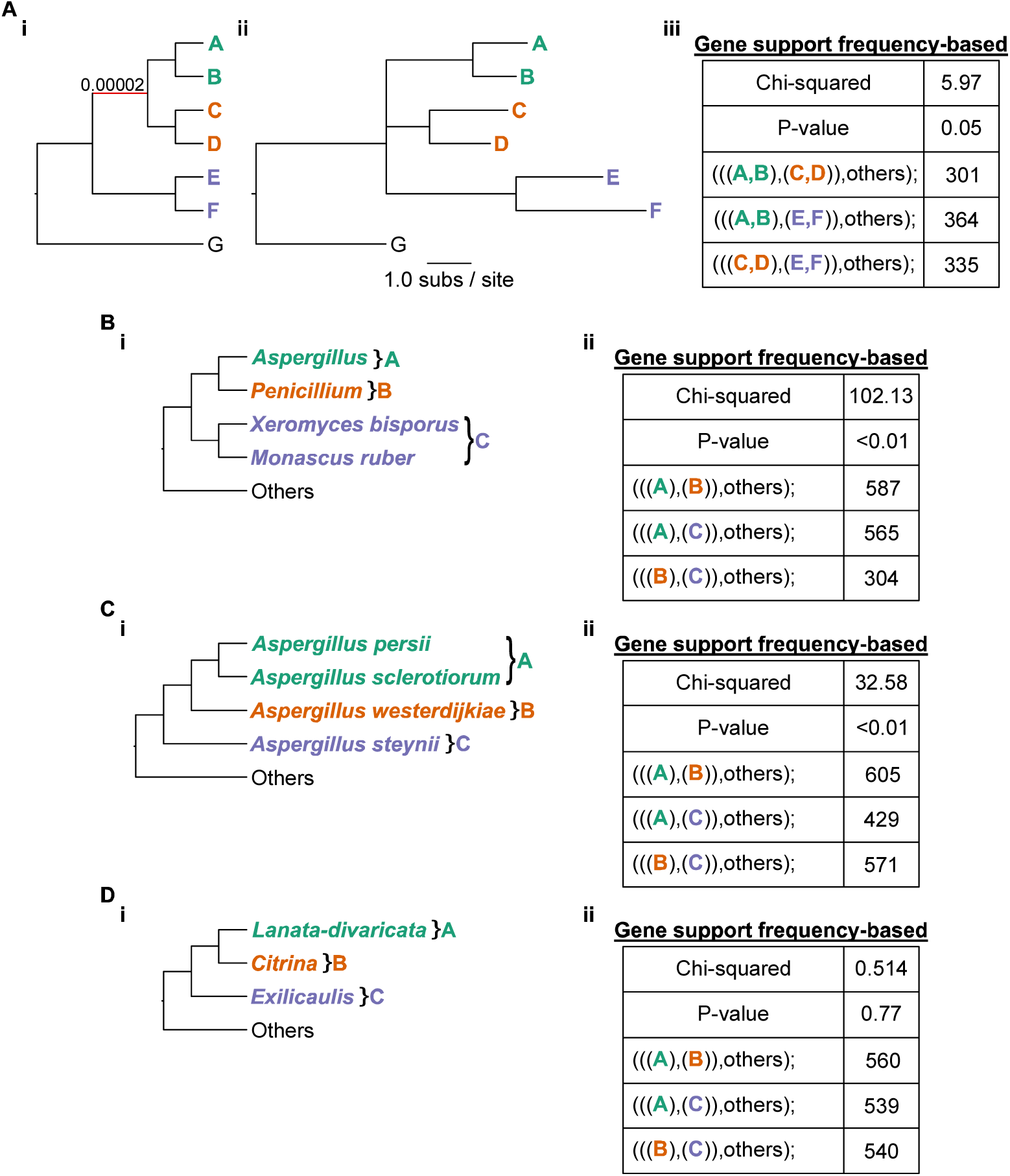
Identifying polytomies from phylogenomic data. (Ai) A cladogram of a simulated species phylogeny with tip names *A-G*. The red branch has a very short branch length of 2×10^−5^ substitutions per site. (Aii) Phylogram of the same phylogeny shows that all other branches are much longer (≥ 1.0 substitutions per site). (Aiii) After reconstructing the evolutionary history from 1,000 alignments simulated from the phylogeny in *Aii*, the hypothesis of a polytomy was tested using gene support frequencies for three alternative rooted topologies defined by the clades of green, orange, and purple taxa. Failure to reject the null hypothesis of equal support among genes for each topology is indicative of a polytomy (χ^2^ = 5.97, p-value = 0.05, Chi-squared test). (B-D) The same approach was then used to examine if there is evidence for a polytomy at three different branches in a phylogeny of filamentous fungi. (D) Support for a polytomy (χ^2^ = 0.514, p-value = 0.77, Chi-squared test) was observed for the relationships between three different sections of *Penicillium* fungi. These results demonstrate the utility of gene-support frequencies for evaluating polytomies and examining incongruence in phylogenomic datasets.

We next examined if there is evidence of polytomies in the evolutionary history of filamentous fungi from the genera *Aspergillus* and *Penicillium*. We examined three branches. The first two branches—one dating back ∼110 million years ago (Figure 3B), and another dating back ∼25 million years ago (Figure 3C)—were not polytomies. In contrast, examination of a ∼60 million-year-old branch involving *Lanata-divaricata, Citrina*, and *Exilicaulis* (Figure 3D), which are major lineages (or sections) in the genus *Penicillium*, was consistent with a polytomy. Given the large number of gene trees used in our analysis (n=1,668), these results are consistent with a rapid radiation or diversification event in the history of *Penicillium* species.

In summary, these results suggest that PhyKIT is useful in identifying polytomies in simulated and empirical datasets. PhyKIT can also be useful for exploring incongruence in phylogenies by calculating gene support frequencies for alternative topologies. Calculations of gene-based support for various topologies can be used in diverse applications, including identifying putative introgression / hybridization events and conducting phylogenetically-based genome-wide association (PhyloGWAS) studies (Pease *et al*. 2016; Steenwyk *et al*. 2019).

## Conclusion

We have developed PhyKIT, a comprehensive toolkit for processing and analyzing MSAs and trees in phylogenomic datasets. PhyKIT is freely available on GitHub (https://github.com/JLSteenwyk/PhyKIT) with extensive documentation and user tutorials (https://jlsteenwyk.com/PhyKIT). PhyKIT is a fast and flexible toolkit for the UNIX shell environment, which allows it to be easily integrated into bioinformatic pipelines. We anticipate PhyKIT will be of interest to biologists from diverse disciplines and with varying degrees of experience in analyzing MSAs and phylogenies. In particular, PhyKIT will likely be helpful in addressing one of the greatest challenges in biology, building, understanding, and deriving meaning from the tree of life.

## Data Availability

All data used will become available in figshare (doi: 10.6084/m9.figshare.13118600) upon publication.

## Acknowledgements

We thank the Rokas lab for helpful discussion and feedback. J.L.S. and A.R. were funded by the Howard Hughes Medical Institute through the James H. Gilliam Fellowships for Advanced Study program. X.X.S. was supported by National Natural Science Foundation of China (No. 32071665) and the Fundamental Research Funds for the Central Universities (No. 2020QNA6019).

## References

Bogomolnaya L. M., R. Pathak, J. Guo, and M. Polymenis, 2006 Roles of the RAM signaling network in cell cycle progression in Saccharomyces cerevisiae. Curr. Genet. 49: 384–92. https://doi.org/10.1007/s00294-006-0069-y

Boiteux S., and S. Jinks-Robertson, 2013 DNA Repair Mechanisms and the Bypass of DNA Damage in Saccharomyces cerevisiae. Genetics 193: 1025–1064. https://doi.org/10.1534/genetics.112.145219

Brunette G. J., M. A. Jamalruddin, R. A. Baldock, N. L. Clark, and K. A. Bernstein, 2019 Evolution-based screening enables genome-wide prioritization and discovery of DNA repair genes. Proc. Natl. Acad. Sci. 116: 19593–19599. https://doi.org/10.1073/pnas.1906559116

Chen M.-Y., D. Liang, and P. Zhang, 2017 Phylogenomic Resolution of the Phylogeny of Laurasiatherian Mammals: Exploring Phylogenetic Signals within Coding and Noncoding Sequences. Genome Biol. Evol. 9: 1998–2012. https://doi.org/10.1093/gbe/evx147

Chikina M., J. D. Robinson, and N. L. Clark, 2016 Hundreds of Genes Experienced Convergent Shifts in Selective Pressure in Marine Mammals. Mol. Biol. Evol. 33: 2182–2192. https://doi.org/10.1093/molbev/msw112

Clark N. L., E. Alani, and C. F. Aquadro, 2012 Evolutionary rate covariation reveals shared functionality and coexpression of genes. Genome Res. 22: 714–720. https://doi.org/10.1101/gr.132647.111

Cock P. J. A., T. Antao, J. T. Chang, B. A. Chapman, C. J. Cox, et al., 2009 Biopython: freely available Python tools for computational molecular biology and bioinformatics. Bioinformatics 25: 1422–1423. https://doi.org/10.1093/bioinformatics/btp163

Degnan J. H., and L. A. Salter, 2005 Gene tree distributions under the coalescent process Evolution (N. Y). 59: 24–37. https://doi.org/10.1111/j.0014-3820.2005.tb00891.x

Doolittle W. F., and E. Bapteste, 2007 Pattern pluralism and the Tree of Life hypothesis. Proc. Natl. Acad. Sci. 104: 2043–2049. https://doi.org/10.1073/pnas.0610699104

Doyle V. P., R. E. Young, G. J. P. Naylor, and J. M. Brown, 2015 Can We Identify Genes with Increased Phylogenetic Reliability? Syst. Biol. 64: 824–837. https://doi.org/10.1093/sysbio/syv041

Felsenstein J., 1978 Cases in which Parsimony or Compatibility Methods will be Positively Misleading. Syst. Biol. 27: 401–410. https://doi.org/10.1093/sysbio/27.4.401

Fourment M., and M. J. Gibbs, 2006 PATRISTIC: a program for calculating patristic distances and graphically comparing the components of genetic change. BMC Evol. Biol. 6: 1. https://doi.org/10.1186/1471-2148-6-1

Hunter J. E., and S. H. Cohen, 2007 Package: igraph. Educ. Psychol. Meas. https://doi.org/10.1177/001316446902900315

Jarvis E. D., S. Mirarab, A. J. Aberer, B. Li, P. Houde, et al., 2014 Whole-genome analyses resolve early branches in the tree of life of modern birds. Science (80-.). 346: 1320–1331. https://doi.org/10.1126/science.1253451

Junier T., and E. M. Zdobnov, 2010 The Newick utilities: high-throughput phylogenetic tree processing in the UNIX shell. Bioinformatics 26: 1669–1670. https://doi.org/10.1093/bioinformatics/btq243

Kapli P., Z. Yang, and M. J. Telford, 2020 Phylogenetic tree building in the genomic age. Nat. Rev. Genet. https://doi.org/10.1038/s41576-020-0233-0

Kück P., and G. C. Longo, 2014 FASconCAT-G: extensive functions for multiple sequence alignment preparations concerning phylogenetic studies. Front. Zool. 11: 81. https://doi.org/10.1186/s12983-014-0081-x

Kumar S., G. Stecher, and K. Tamura, 2016 MEGA7: Molecular Evolutionary Genetics Analysis Version 7.0 for Bigger Datasets. Mol. Biol. Evol. https://doi.org/10.1093/molbev/msw054

Lake J. A., 1991 The order of sequence alignment can bias the selection of tree topology. Mol. Biol. Evol. https://doi.org/10.1093/oxfordjournals.molbev.a040654

Lanyon S. M., 1988 The Stochastic Mode of Molecular Evolution: What Consequences for Systematic Investigations? Auk 105: 565–573. https://doi.org/10.1093/auk/105.3.565

Laumer C. E., R. Fernández, S. Lemer, D. Combosch, K. M. Kocot, et al., 2019 Revisiting metazoan phylogeny with genomic sampling of all phyla. Proc. R. Soc. B Biol. Sci. 286: 20190831. https://doi.org/10.1098/rspb.2019.0831

Li Y., J. L. Steenwyk, Y. Chang, Y. Wang, T. Y. James, et al., 2020 A genome-scale phylogeny of Fungi; insights into early evolution, radiations, and the relationship between taxonomy and phylogeny. bioRxiv 2020.08.23.262857. https://doi.org/10.1101/2020.08.23.262857

Liu L., J. Zhang, F. E. Rheindt, F. Lei, Y. Qu, et al., 2017 Genomic evidence reveals a radiation of placental mammals uninterrupted by the KPg boundary. Proc. Natl. Acad. Sci. 114: E7282–E7290. https://doi.org/10.1073/pnas.1616744114

Lujan S. A., J. S. Williams, Z. F. Pursell, A. A. Abdulovic-Cui, A. B. Clark, et al., 2012 Mismatch Repair Balances Leading and Lagging Strand DNA Replication Fidelity, (C. E. Pearson, Ed.). PLoS Genet. 8: e1003016. https://doi.org/10.1371/journal.pgen.1003016

Novick P., and R. Schekman, 1979 Secretion and cell-surface growth are blocked in a temperature-sensitive mutant of Saccharomyces cerevisiae. Proc. Natl. Acad. Sci. U. S. A. 76: 1858–62. https://doi.org/10.1073/pnas.76.4.1858

One Thousand Plant Transcriptomes Initiative, 2019 One thousand plant transcriptomes and the phylogenomics of green plants. Nature 574: 679–685. https://doi.org/10.1038/s41586-019-1693-2

Pazos F., and A. Valencia, 2001 Similarity of phylogenetic trees as indicator of protein–protein interaction. Protein Eng. Des. Sel. 14: 609–614. https://doi.org/10.1093/protein/14.9.609

Pease J. B., D. C. Haak, M. W. Hahn, and L. C. Moyle, 2016 Phylogenomics Reveals Three Sources of Adaptive Variation during a Rapid Radiation, (D. Penny, Ed.). PLOS Biol. 14: e1002379. https://doi.org/10.1371/journal.pbio.1002379

Philippe H., H. Brinkmann, D. V. Lavrov, D. T. J. Littlewood, M. Manuel, et al., 2011 Resolving Difficult Phylogenetic Questions: Why More Sequences Are Not Enough, (D. Penny, Ed.). PLoS Biol. 9: e1000602. https://doi.org/10.1371/journal.pbio.1000602

Phillips M. J., and D. Penny, 2003 The root of the mammalian tree inferred from whole mitochondrial genomes. Mol. Phylogenet. Evol. 28: 171–185. https://doi.org/10.1016/S1055-7903(03)00057-5

Revell L. J., 2012 phytools: an R package for phylogenetic comparative biology (and other things). Methods Ecol. Evol. 3: 217–223. https://doi.org/10.1111/j.2041-210X.2011.00169.x

Robinson D. F., and L. R. Foulds, 1981 Comparison of phylogenetic trees. Math. Biosci. 53: 131–147. https://doi.org/10.1016/0025-5564(81)90043-2

Rokas A., and S. B. Carroll, 2006 Bushes in the Tree of Life. PLoS Biol. 4: e352. https://doi.org/10.1371/journal.pbio.0040352

Salichos L., and A. Rokas, 2013 Inferring ancient divergences requires genes with strong phylogenetic signals. Nature 497: 327–331. https://doi.org/10.1038/nature12130

Sang T., and Y. Zhong, 2000 Testing Hybridization Hypotheses Based on Incongruent Gene Trees, (R. Olmstead, Ed.). Syst. Biol. 49: 422–434. https://doi.org/10.1080/10635159950127321

Sato T., Y. Yamanishi, M. Kanehisa, and H. Toh, 2005 The inference of protein-protein interactions by co-evolutionary analysis is improved by excluding the information about the phylogenetic relationships. Bioinformatics 21: 3482–3489. https://doi.org/10.1093/bioinformatics/bti564

Sayyari E., and S. Mirarab, 2018 Testing for Polytomies in Phylogenetic Species Trees Using Quartet Frequencies. Genes (Basel). 9. https://doi.org/10.3390/genes9030132

Shen X.-X., L. Salichos, and A. Rokas, 2016a A Genome-Scale Investigation of How Sequence, Function, and Tree-Based Gene Properties Influence Phylogenetic Inference. Genome Biol. Evol. 8: 2565–2580. https://doi.org/10.1093/gbe/evw179

Shen X.-X., X. Zhou, J. Kominek, C. P. Kurtzman, C. T. Hittinger, et al., 2016b Reconstructing the Backbone of the Saccharomycotina Yeast Phylogeny Using Genome-Scale Data. G3 Genes|Genomes|Genetics 6: 3927–3939. https://doi.org/10.1534/g3.116.034744

Shen X.-X., D. A. Opulente, J. Kominek, X. Zhou, J. L. Steenwyk, et al., 2018 Tempo and Mode of Genome Evolution in the Budding Yeast Subphylum. Cell 175: 1533-1545.e20. https://doi.org/10.1016/j.cell.2018.10.023

Shen X.-X., J. L. Steenwyk, A. L. Labella, D. A. Opulente, X. Zhou, et al., 2020 Genome-scale phylogeny and contrasting modes of genome evolution in the fungal phylum Ascomycota. bioRxiv. https://doi.org/10.1101/2020.05.11.088658

Smith S. A., J. W. Brown, and J. F. Walker, 2018 So many genes, so little time: A practical approach to divergence-time estimation in the genomic era, (H. Escriva, Ed.). PLoS One 13: e0197433. https://doi.org/10.1371/journal.pone.0197433

Song Y., S. A. Cheon, K. E. Lee, S.-Y. Lee, B.-K. Lee, et al., 2008 Role of the RAM network in cell polarity and hyphal morphogenesis in Candida albicans. Mol. Biol. Cell 19: 5456–77. https://doi.org/10.1091/mbc.e08-03-0272

Steenwyk J. L., X.-X. Shen, A. L. Lind, G. H. Goldman, and A. Rokas, 2019 A Robust Phylogenomic Time Tree for Biotechnologically and Medically Important Fungi in the Genera Aspergillus and Penicillium, (J. P. Boyle, Ed.). MBio 10. https://doi.org/10.1128/mBio.00925-19

Steenwyk J. L., T. J. Buida, Y. Li, X.-X. Shen, and A. Rokas, 2020 ClipKIT: a multiple sequence alignment-trimming algorithm for accurate phylogenomic inference. bioRxiv 2020.06.08.140384. https://doi.org/10.1101/2020.06.08.140384

Struck T. H., 2014 TreSpEx–-Detection of Misleading Signal in Phylogenetic Reconstructions Based on Tree Information. Evol. Bioinforma. 10: EBO.S14239. https://doi.org/10.4137/EBO.S14239

Sundin L. J. R., G. J. Guimaraes, and J. G. Deluca, 2011 The NDC80 complex proteins Nuf2 and Hec1 make distinct contributions to kinetochore-microtubule attachment in mitosis. Mol. Biol. Cell 22: 759–68. https://doi.org/10.1091/mbc.E10-08-0671

Talevich E., B. M. Invergo, P. J. Cock, and B. A. Chapman, 2012 Bio.Phylo: A unified toolkit for processing, analyzing and visualizing phylogenetic trees in Biopython. BMC Bioinformatics 13: 209. https://doi.org/10.1186/1471-2105-13-209

Tsubouchi H., and H. Ogawa, 2000 Exo1 Roles for Repair of DNA Double-Strand Breaks and Meiotic Crossing Over in Saccharomyces cerevisiae, (T. D. Fox, Ed.). Mol. Biol. Cell 11: 2221–2233. https://doi.org/10.1091/mbc.11.7.2221

Virtanen P., R. Gommers, T. E. Oliphant, M. Haberland, T. Reddy, et al., 2020 SciPy 1.0: fundamental algorithms for scientific computing in Python. Nat. Methods. https://doi.org/10.1038/s41592-019-0686-2

Walker J. F., N. Walker-Hale, O. M. Vargas, D. A. Larson, and G. W. Stull, 2019 Characterizing gene tree conflict in plastome-inferred phylogenies. PeerJ 7: e7747. https://doi.org/10.7717/peerj.7747

Walsh H. E., M. G. Kidd, T. Moum, and V. L. Friesen, 1999 Polytomies and the Power of Phylogenetic Inference. Evolution (N. Y). 53: 932. https://doi.org/10.2307/2640732

Weigert A., C. Helm, M. Meyer, B. Nickel, D. Arendt, et al., 2014 Illuminating the Base of the Annelid Tree Using Transcriptomics. Mol. Biol. Evol. 31: 1391–1401. https://doi.org/10.1093/molbev/msu080

Wolfe N. W., and N. L. Clark, 2015 ERC analysis: web-based inference of gene function via evolutionary rate covariation: Fig. 1. Bioinformatics btv454. https://doi.org/10.1093/bioinformatics/btv454

